# Endothelial Cell Cycle State Determines Propensity for Arterial-Venous Fate

**DOI:** 10.1101/2020.08.12.246512

**Authors:** Nicholas W. Chavkin, Gael Genet, Mathilde Poulet, Nafiisha Genet, Corina Marziano, Hema Vasavada, Elizabeth A. Nelson, Anupreet Kour, Stephanie P. McDonnell, Mahalia Huba, Kenneth Walsh, Karen K. Hirschi

## Abstract

Formation and maturation of a functional blood vascular system is required for the development and maintenance of all tissues in the body. During the process of blood vessel development, primordial endothelial cells are formed and become specified toward arterial or venous fates to generate a circulatory network that provides nutrients and oxygen to, and removes metabolic waste from, all tissues^1-3^. Specification of arterial and venous endothelial cells occurs in conjunction with suppression of endothelial cell cycle progression^4,5^, and endothelial cell hyperproliferation is associated with potentially lethal arterial-venous malformations^6^. However, the mechanistic role that cell cycle state plays in arterial-venous specification is unknown. Herein, studying retinal vascular development in Fucci2aR reporter mice^7^, we found that venous and arterial endothelial cells are in distinct cell cycle states during development and in adulthood. That is, venous endothelial cells reside in early G1 state, while arterial endothelial cells reside in late G1 state. Endothelial cells in early vs. late G1 exhibited significant differences in gene expression and activity, especially among BMP/TGF-β signaling components. The early G1 state was found to be essential for BMP4-induced venous specification, whereas late G1 state is essential for TGF-β1-induced arterial specification. In a mouse model of endothelial cell hyperproliferation and disrupted vascular remodeling, pharmacological inhibition of endothelial cell cycle rescues the arterial-venous specification defects. Collectively, our results show that endothelial cell cycle control plays a key role in arterial-venous network formation, and distinct cell cycle states provide distinct windows of opportunity for the molecular induction of arterial vs. venous specification.

## Introduction

Healthy tissue development and maintenance requires a functional blood circulatory network comprised of arterial and venous blood vessels lined with specialized endothelial cells. Acquisition of these specialized arterial and venous endothelial cell phenotypes generally occurs in conjunction with suppression of endothelial cell cycle progression^1,4,5^. However, we lack understanding of mechanisms that coordinately regulate endothelial cell growth suppression and phenotypic specialization during vascular remodeling, which creates significant roadblocks for clinical therapies, tissue engineering and regenerative medicine.

Our previous work has shown that shear stress, specifically at magnitudes typically found in arteries and arterioles, activates a Notch-Cx37-p27 signaling axis to promote endothelial cell cycle arrest, and that this enables the upregulation of arterial genes^4^. However, it is not clear whether a specific state of the cell cycle plays a role in venous endothelial cell specification, or whether distinct cell cycle states control the differential specification of arterial and venous endothelial cells. In this regard, distinct signaling pathways have been implicated in the upregulation of arterial or venous genes, including TGF-β and BMP^8-12^, respectively, but how these signaling pathways function in coordination with cell cycle state to induce specific endothelial cell phenotypes is also not known.

To fill these knowledge gaps, we created mice expressing the Fluorescent Ubiquitination Cell Cycle Indicator (FUCCI) reporter specifically in endothelial cells. Using these mice, we demonstrated that endothelial cells in veins/venules vs. arteries/arterioles are in distinct cell cycle states during vascular development and in adulthood; early G1 vs. late G1, respectively. Although both early G1 and late G1 represent states of “growth arrest”, in embryonic stem cells, these states are molecularly distinct and represent distinct windows of opportunity for the induction of mesoderm/endoderm vs. ectoderm lineages^15,16^.

We then performed studies using human umbilical vein endothelial cells transduced with a lentivirus expressing the FUCCI reporter^13^ (HUVEC-FUCCI) to demonstrate that shear stress typical of veins/venules (4 dynes/cm^2^) promotes early G1 arrest; whereas, shear stress typical of arteries/arterioles (12 dynes/cm^2^) promotes late G1 arrest. Furthermore, these different endothelial cell cycle states are shown to provide distinct windows of opportunity for gene expression in response to extrinsic signals. That is, components of the BMP/TGF-β signaling pathways are shown to be differentially regulated in early vs. late G1, and BMP signaling induces venous gene expression only in early G1; whereas, TGF-β induces arterial gene expression only in late G1. Finally, using Cx37-deficient mice that exhibit endothelial cell hyperproliferation, dysregulated vascular remodeling and impaired arterial development, we showed that pharmacological induction of endothelial cell cycle arrest in late G1 state rescues these vascular defects and restores normal arterial-venous network formation.

These studies reveal a critical and previously unknown molecular connection between endothelial cell cycle state and fate; specifically, endothelial cell cycle state determines the propensity for arterial vs. venous fate specification.

## Results

### Arterial-Venous Endothelial Cell Cycle State

To determine the cell cycle state of endothelial cells during arterial-venous specification, we used mice expressing the Fluorescent Ubiquitination Cell Cycle Indicator (FUCCI) reporter, which enables clear distinction among cells in early G1, late G1 and S/G2/M states (**Fig. 1A**). To specifically label endothelial cells, we crossed mice expressing a FUCCI reporter with a flox-stop-flox cassette^7^ with mice expressing the endothelial-specific Cdh5-CreER^T2,14^. In these tamoxifen-treated mice, at postnatal day (P)6, we examined retinal endothelial cells in the developing arterial-venous network (**Fig. 1B**). We found that endothelial cells in S/G2/M states (green) are in the remodeling areas closer to venous vessels. Endothelial cells in and near the arterial branches are in late G1 state (red) (**Fig. 1C**); whereas, endothelial cells in and near the venous branches are in early G1 (unlabeled) (**Fig. 1D**). These patterns persisted in P15 retinal vasculature, in which arterial and venous branches have matured (**Fig. 1E-G**), and into adulthood (**Ext. Fig. 1A-C**).

**Figure 1.**
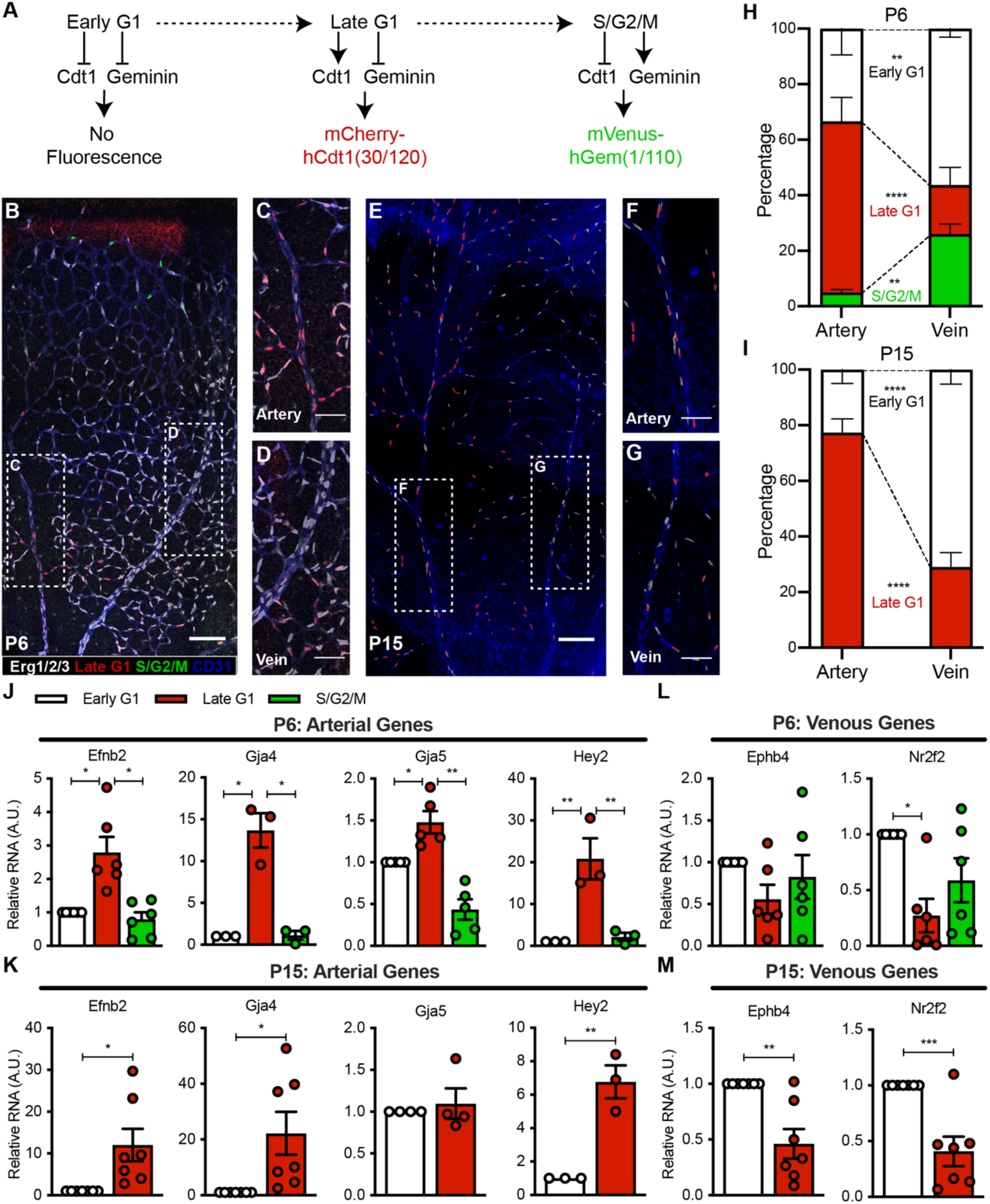
Endothelial Cell Cycle State during Retina Vascular Development. **A**) FUCCI reporter distinguishes early G1, late G1 and S/G2/M cell cycle states. **B**) P6 retinas of Fucci2aR mice imaged for CD31, Erg1/2/3, hCdt1(30/120) and hGem(1/110) (scale bar = 200μm), magnified on **C**) artery and **D**) vein (scale bar = 50μm). **E**) P15 retinas imaged, magnified on **F**) artery and **G**) vein. **H-I**) Cell cycle states quantified in arteries and veins at P6 and P15. Gene expression of retina endothelial cells in different cell cycle states at P6 and P15 quantified for **J-K**) arterial genes and **L-M**) venous genes.

Quantification of endothelial cell cycle states in blood vessels at P6 and P15 confirmed that arterial vessels have a greater percentage of endothelial cells in late G1, while venous vessels have a greater percentage of endothelial cells in early G1 (**Fig. 1H-I**). Additionally, we found that throughout the course of the retinal vascular plexus maturation, early G1 endothelial cells associate closer to veins/venules while late G1 endothelial cells associate closer to arteries/arterioles (**Ext. Fig. 1D-E**). Consistent with these findings, we observed that arterial shear flow forces (12 dynes/cm^2^) increase the proportion of HUVEC-FUCCI in late G1 state and concomitantly reduce the proportion in early G1 state, compared to venous shear flow forces (4 dynes/cm^2^), which promote early G1 state (**Ext. Fig. 1F**), suggesting that differential shear flow forces can mediate endothelial cell cycle state.

To further investigate the phenotypes of endothelial cells in distinct cell cycle states, we used fluorescence activated cell sorting (FACS) to isolate P6 and P15 retinal CD31+CD45-endothelial cells from our FUCCI reporter mice (sorting strategy shown in **Fig. 2C**). The endothelial cells in early G1, late G1 and S/G2/M states were processed for RNA isolation and qPCR analysis. We found that endothelial cells in late G1 exhibit significantly higher expression of arterial genes (**Fig. 1J-K; Ext. Fig. 1G-H**), and cells in early G1 exhibit significantly higher expression of venous genes (**Fig. 1L-M**). Collectively, these results revealed that endothelial cells in venous vs. arterial branches exhibit distinct cell cycle states; early G1 vs. late G1.

**Figure 2.**
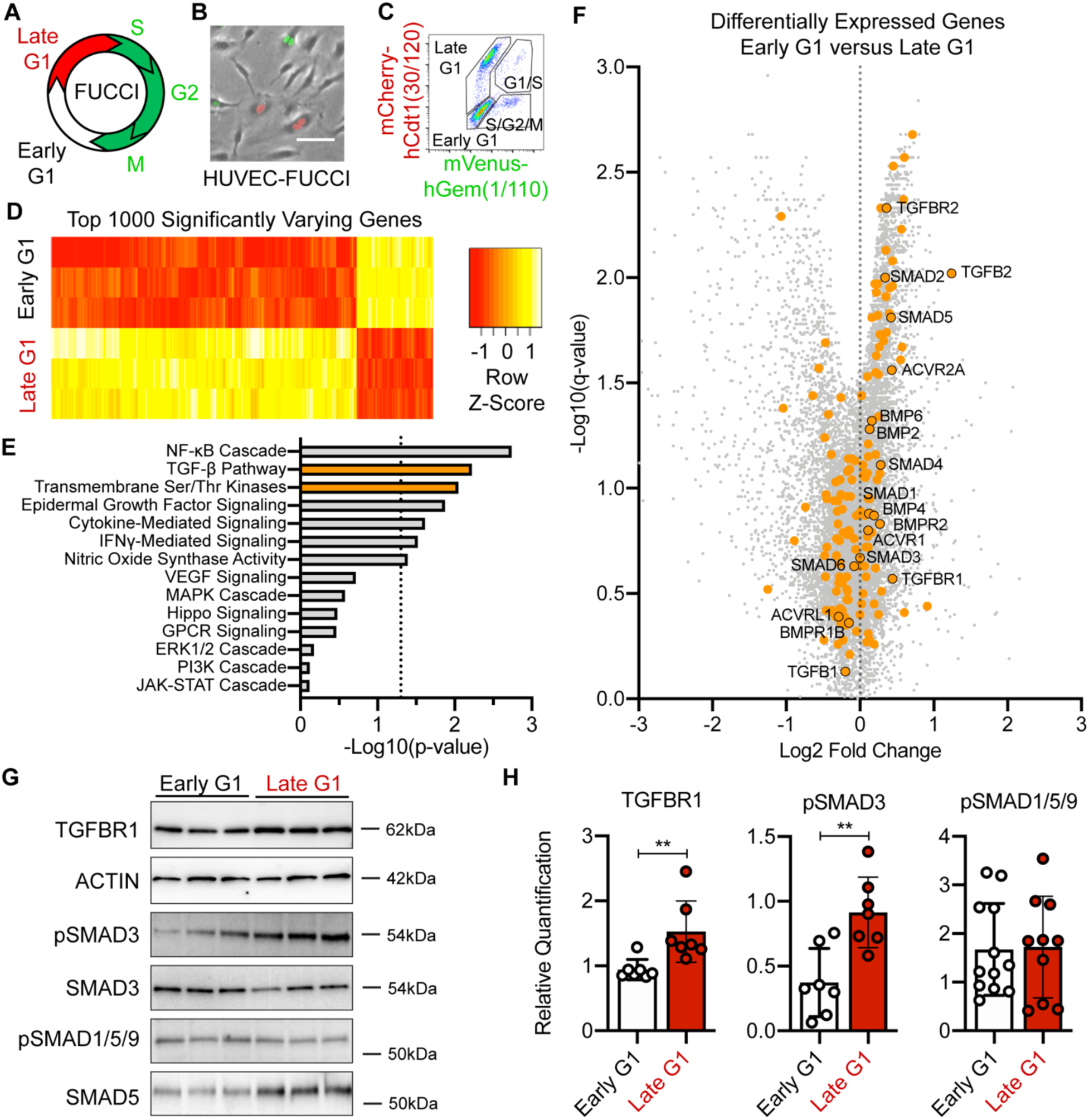
Endothelial Cell Cycle-Dependent Regulation of TGF-β/BMP Pathway. **A**) HUVEC-FUCCI reporter distinguishes early G1, late G1 and S/G2/M cell cycle states, visualized by **B**) fluorescent imaging, and **C**) FACS. In bulk RNA sequencing of early G1 and late G1 HUVEC-FUCCI, **D**) top 1000 significantly varying genes, **E**) Signaling Pathway GO Term analysis, and **F**) Volcano plot of fold-change against the log10(q-value) (TGF-β/BMP signaling pathways in orange). **G**) Western blots of TGF-β/BMP signaling proteins in early G1 and late G1, quantified in **H**).

### Cell Cycle Regulation of TGF-β/BMP Signaling

Recently, embryonic stem cell differentiation towards specific lineages was found to be controlled by cell cycle state regulation of gene expression, chromatin remodeling and transcription factor binding^15,16^. To begin to investigate the mechanistic role of cell cycle state in endothelial cells, we performed gene expression analyses of endothelial cells in distinct cell cycle states. To do so, we FACS-isolated HUVEC-FUCCI cells into early G1, late G1 and S/G2/M states (**Fig. 2A-C**) and isolated RNA therefrom.

Bulk RNA sequencing analysis of endothelial cells in different cell cycle states revealed high transcriptional variation between early G1 and late G1 states (**Fig. 2D**). We performed Gene Ontology analysis to determine which signaling pathways were significantly different and found that the TGF-β and transmembrane serine/threonine kinase signaling pathways are significantly variable between early G1 and late G1 (**Fig. 2E**). These two pathways contain many genes in the TGF-β and BMP signaling pathways, which are known to be involved in arterial-venous specification^11,12,17,18^, and we found many of these genes differentially regulated in early vs. late G1 (**Fig. 1F**). Furthermore, via Western Blot analysis of endothelial cells in early G1 and late G1, we also found that the protein expression of some TGF-β signaling components (phospho-SMAD3 and TGFBR1) are increased in late G1 (**Fig. 2G-H, Ext. Fig. 2A-B**), suggesting that TGF-β signaling may be more active in late G1 and less active in early G1. We found no differences in noncanonical TGF-β signaling through ERK1/2 or AKT between cell cycle states or after TGF-β1/BMP4 stimulation (**Ext. Fig. 2C-F**). Thus, these results suggest that canonical SMAD-mediated TGF-β and BMP signaling may be differentially regulated in distinct endothelial cell cycle states, which could then enable arterial vs. venous gene induction after TGF-β or BMP stimulation, respectively.

### Cell Cycle Regulation of Arterial-Venous Fate

We then examined whether TGF-β and BMP signaling are activated in endothelial cells in distinct cell cycle states. The key signaling proteins for the TGF-β/BMP signaling pathways are SMAD proteins^19^. SMAD4 serves as the intermediate co-activator, where TGF-β1 induces SMAD2/3 binding to SMAD4 and BMP4 induces SMAD1/5 binding to SMAD4^20^, and these transcription factor complexes function to promote gene expression (**Fig. 3A**). Using our HUVEC-FUCCI, we FACS-isolated endothelial cells in early G1 and late G1 states, and performed SMAD4 co-immunoprecipitation. We found that TGF-β1 (1 ng/ml for 2 hr) induces greater SMAD2/3 binding to SMAD4 in late G1 state, while BMP4 (5 ng/ml for 2 hr) induces greater SMAD1/5 binding to SMAD4 in early G1 state (**Fig. 3B-C**). Thus, BMP and TGF-β signaling appear to be differentially active in early G1 vs. late G1, respectively.

**Figure 3.**
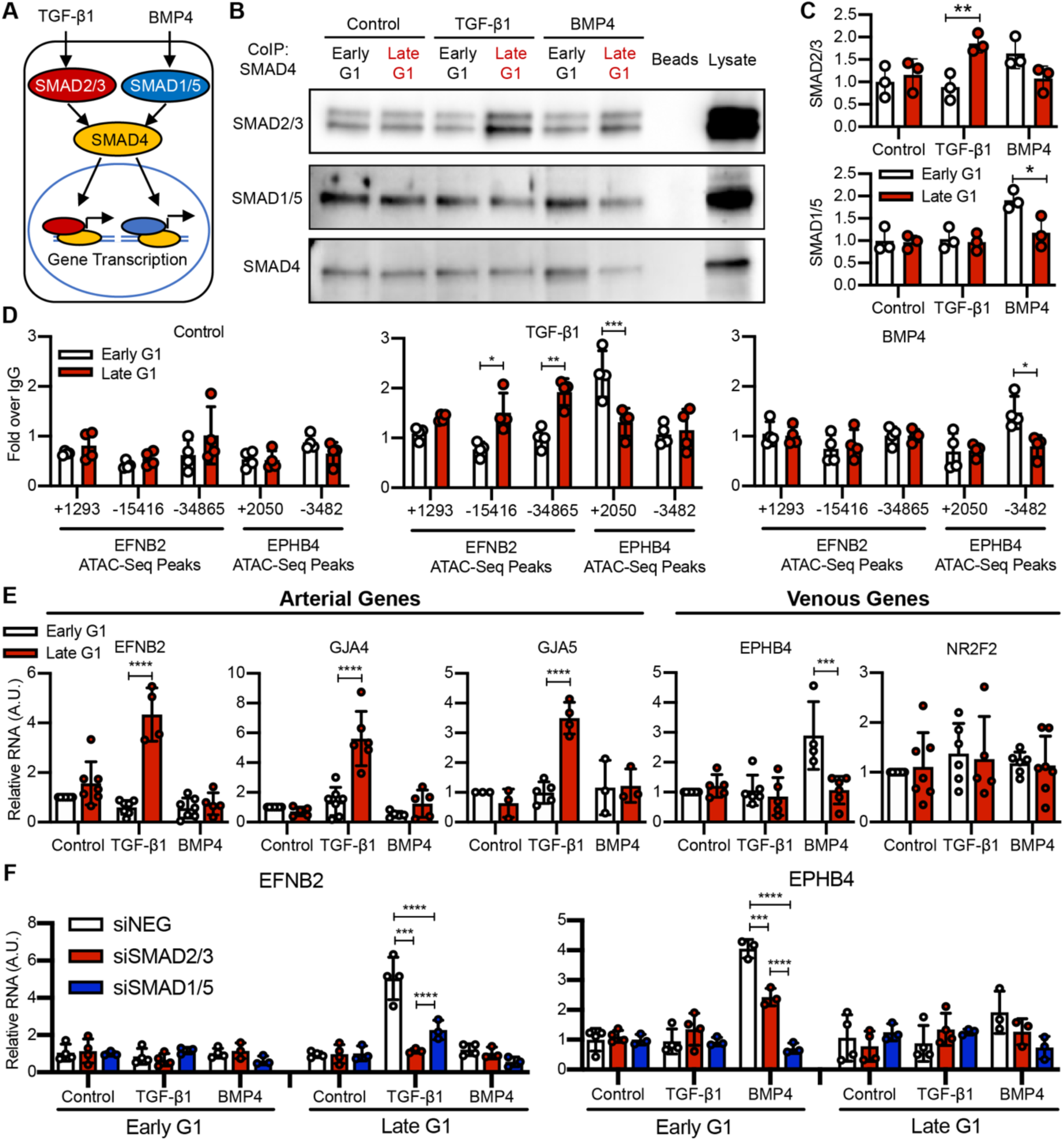
Endothelial Cell Cycle-Dependent Arterial-venous Specification via TGF-β/BMP Signaling. **A**) Overview of the TGF-β/BMP signaling pathway. **B**) Western blot for SMAD proteins of lysates from SMAD4 co-immunoprecipitation after TGF-β1/BMP4-treated early G1 and late G1 HUVEC-FUCCI, quantified in **C**). **D**) qRT-PCR analysis of DNA regions near EFNB2 and EPHB4 binding to SMAD4 complexes by chromatin-immunoprecipitation. **E**) TGF-β1/BMP4 induction of arterial and venous genes in early G1 and late G1 HUVEC-FUCCI. **F**) TGF-β1/BMP4 induction of EFNB2 and EPHB4 after SMAD2/3 or SMAD1/5 siRNA knockdown.

To investigate whether their downstream target genes were differentially susceptible to SMAD4 transcription factor binding in endothelial cells in distinct cell cycle states, we first performed ATAC-Sequencing to identify regions of DNA near arterial-venous genes that were open in HUVEC-FUCCI in early G1 vs. late G1 states (**Ext. Fig. 3A-B**). Surprisingly, we found no significant differences in the size or location of the open chromatin regions near known arterial-venous genes (EFNB2 and EPHB4, respectively). We then performed SMAD4 chromatin immunoprecipitation PCR to quantify SMAD4 transcription factor binding to their specific ATAC-seq peaks in HUVEC-FUCCI in early G1 and late G1 states, in response to TGF-β or BMP stimulation. We found that TGF-β1 (1 ng/ml for 4 hr) induces more SMAD4 binding to ATAC-Seq peaks near the EFNB2 gene, while BMP4 (5 ng/ml for 4 hr) induces more SMAD4 binding to ATAC-Seq peaks near the EPHB4 gene (**Fig. 3D**).

To further test whether BMP4 and TGF-β1 signaling promote venous vs. arterial gene expression in distinct cell cycle states, we treated HUVEC-FUCCI cells, either non-sorted (control) or FACS-isolated in early G1 and late G1 states, with BMP4 or TGF-β1 for 8 hr, and measured changes in mRNA expression via qPCR. Importantly, endothelial cell cycle state was not changed within this treatment duration (**Ext. Fig. 3C).** We found that TGF-β1 does not induce venous gene expression in any condition, and induces arterial genes only in endothelial cells in late G1 state. Conversely, BMP4 induces only venous gene EPHB4 and only in endothelial cells in early G1 (**Fig. 3E**). Thus, we found that cell cycle state-mediated activation of the TGF-β and BMP signaling pathways enables differential regulation of arterial and venous genes, respectively.

Finally, we performed knockdown experiments of SMAD genes to determine the requirement for SMAD signaling in TGF-β1- and BMP4-induced arterial-venous gene expression. Transfection of siRNA yielded >80% knockdown of targeted SMAD genes (**Ext. Fig. 3D**). We found that siRNA-mediated knockdown of SMAD2/3 prevents TGF-β1-induced EFNB2 gene induction in late G1, and knockdown of SMAD1/5 prevents BMP4-induced EPHB4 gene induction in early G1 (**Fig. 3F**). These results suggest that late G1 state is required for TGF-β1-induced arterial gene induction through SMAD2/3, and early G1 state is required for BMP4-induced venous gene induction through SMAD1/5 in endothelial cells.

### Rescue of Arterial-Venous Fate Defects

To investigate the role of endothelial cell cycle state in arterial-venous specification in vivo, we tested whether pharmacological manipulation of cell cycle state could rescue arterial-venous specification defects associated with endothelial cell hyperproliferation. We used Connexin (Cx)37-deficient mice (Cx37-KO) in which we previously found endothelial cell hyperproliferation, downregulated cell cycle inhibitor p27 (Cdkn1b), and impaired arterial blood vessel maturation^4^. p27 induces G1 arrest in cells by interacting and inhibiting the Cyclin D-CDK4 and Cyclin E-CDK2 complexes^21,22^, which normally function to promote G1-to-S transition^23^.

Cell cycle state can be regulated through pharmacological inhibition of CDK proteins. Specifically, we found that CDK4/6i (Palbociclib, PD-0332991) reduces active cycling (S/G2/M) and promotes late G1 arrest in endothelial cells (**Ext. Fig. 4A**). Therefore, we treated Cx37-KO mice with CDK4/6i during retinal vascular development and assessed its impact on vascular remodeling and arterial-venous specification. We found that the vascular hyper-density and arterial maturation defects (indicated by reduced αSMA coverage) observed in P6 retinas of Cx37-KO mice are both rescued with CDK4/6i treatment (**Fig. 4A-E**). Importantly, we found that CDK4/6i treatment does not significantly affect mouse growth/weight (**Ext. Fig. 4B**).

**Figure 4.**
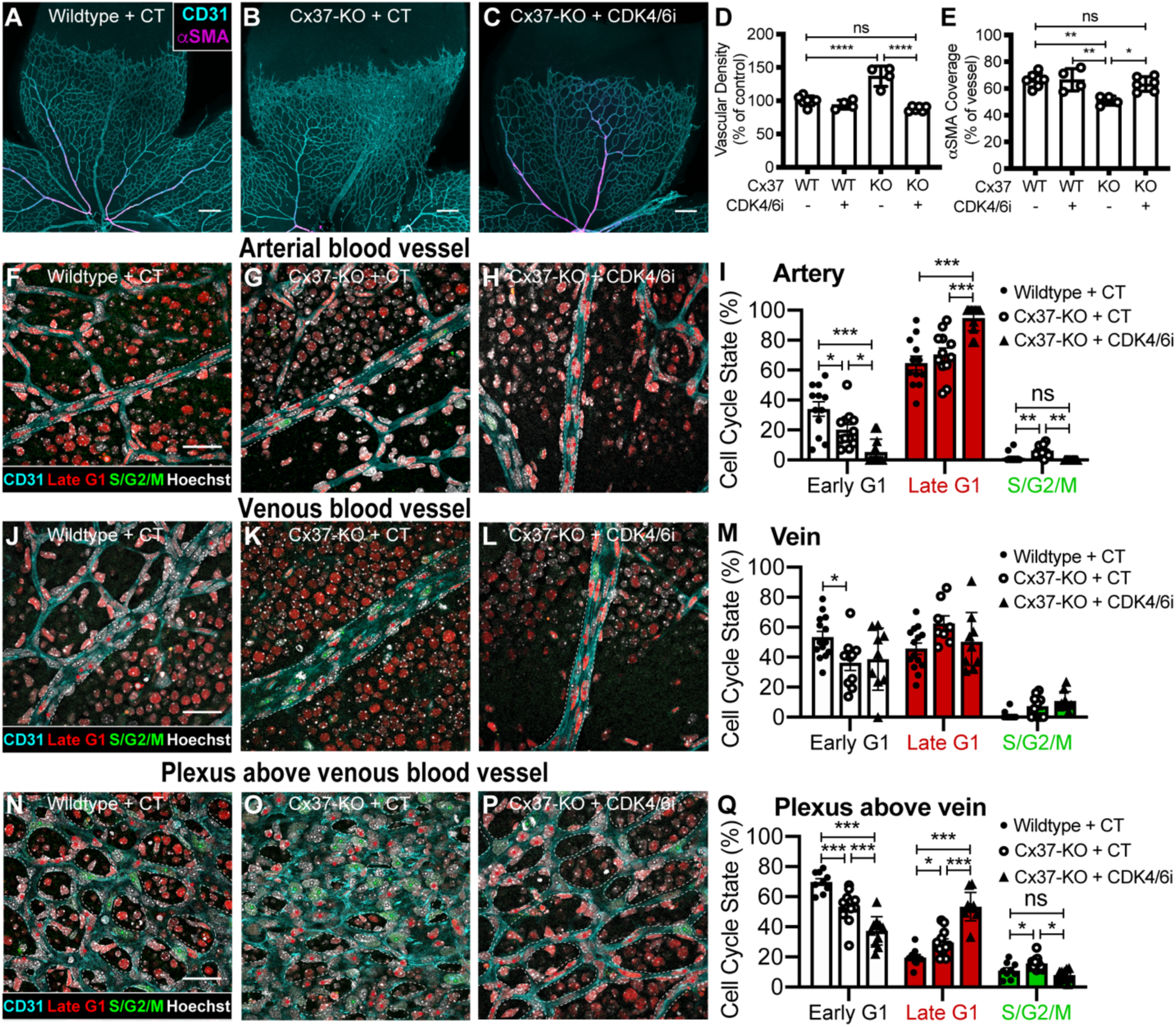
Rescue of Arterial-venous Development Defects with Pharmacological CDK4/6 Inhibition. **A-C**) P6 retinal vasculature of WT, Cx37-KO and Cx37-KO+CDK4/6i treated mice imaged for CD31 and αSMA (scale bars = 200μm), quantified for **D**) vascular density, and **E**) αSMA coverage. P6 retinal vasculature of Fucci2, Fucci2+Cx37-KO and Fucci2+Cx37-KO+CDK4/6i treatment imaged for CD31, hCdt1(30/120), hGem(1/110) and Erg1/2/3 and quantified for cell cycle state in **F-I**) arterial blood vessels, **J-M**) venous blood vessels, and **N-Q**) plexi above venous blood vessels (scale bars = 50μm, vessels outlined in dotted white lines, cell cycle state highlighted with colored stars).

We then investigated developing retinal endothelial cell cycle state in Cx37^-/-^;R26p-FUCCI2 mice compared to wildtype controls, as well as the effects of CDK4/6i treatment on endothelial cell cycle state during vascular remodeling. Firstly, we found that, in Cx37-KO mice compared to wildtype controls, a higher proportion of endothelial cells in arterial branches and plexi about the venous branches are actively cycling (S/G2/M), and a lower proportion of endothelial cells in arterial and venous branches and their associated plexi are in early G1 (**Fig. 4F-Q, Ext. Fig. 4C-F**). Interestingly, a higher proportion of endothelial cells in the plexi above the venous and arterial branches are also in late G1 (**Fig. 1N-Q and Ext. Fig. 4C-F**).

When Cx37^-/-^;R26p-FUCCI2 mice were treated with the CDK4/6i, they exhibited an increased proportion of endothelial cells in arterial branches in late G1 state and a reduced proportion in early G1 and S/G2/M cell cycle states (**Fig. 4F-I**). We did not observe large changes in the cell cycle state of endothelial cells within venous branches themselves or the plexi above the arterial blood vessels (**Fig. 4J-M, Ext. Ext. Fig. 4C-F**). However, in the remodeling plexi above the venous blood vessels, we found significantly less endothelial cells in early G1 and S/G2/M states and significantly more in late G1 after CDK4/6i treatment (**Fig. 4N-Q**). We further confirmed that CDK4/6i treatment inhibits cell proliferation assessed by EdU incorporation (**Ext. Fig. 4G-J**). Thus, we found that inducing late G1 arrest in retinal endothelial cells in the Cx37-KO mice is sufficient to rescue their defects in arterial development and vascular remodeling.

## Discussion

These studies reveal that arterial and venous endothelial cells reside in distinct cell cycle states during development and into adulthood. Although both venous and arterial endothelial cells are “growth arrested”, endothelial cells in venous branches are in an early G1 phase and endothelial cells in arterial branches are in a late G1 phase of cell cycle. Notably, the magnitude of shear stress that endothelial cells experience was shown to control the cell cycle state in which they reside. Specifically, shear stress magnitudes typically experienced by venous endothelial cells promote early G1 arrest, whereas the greater blood flow forces sensed by arterial endothelial cells promote late G1 arrest. These effects are likely mediated via the activation of distinct downstream signaling pathways that differentially regulate cell cycle state, consistent with previous studies that showed arterial shear stress activates Notch signaling to promote late G1 arrest that enables arterial gene expression^4^.

Our studies are the first to identify a molecular link between cell cycle state and endothelial cell fate, and they provide a framework that integrates other observations in the field. For example, although BMP^12^ and TGF-β^10^ signaling had been shown to promote venous and arterial gene expression, respectively, it was not clear how these pathways are coordinated with the shear stress differences that are associated with venous vs. arterial systems. Our findings show that the early G1 state, induced by low shear stress, will license endothelial cells to be permissive for BMP signaling and thereby allow venous specification. In contrast, high shear stress promotes a late G1 state, and this is more permissive for TGF-β signaling that promotes arterial specification. Thus, these studies provide a paradigm of arterial vs. venous phenotypic specialization that involves flow-mediated control of distinct signaling pathways that serve to position endothelial cells in different cell cycle states and thereby enable arterial vs. venous patterns of gene expression.

Other recent studies have highlighted the importance of cell cycle regulation in endothelial cell differentiation. Coronary artery development is impaired when cell cycle control is disrupted in the progenitor endothelial cells coming from the sinus venosus^5^. Also, p27-mediated cell cycle arrest is required for endothelial cells to undergo hemogenic specification during definitive hematopoiesis^26^. Thus, regulation of cell cycle state may be required for the phenotypic specialization of all endothelial cell subtypes. Further, given that some endothelial cells can undergo phenotypic specification in the absence of blood flow^24,25^, it is possible that control of cell cycle by other microenvironment factors enables phenotypic specialization under these unique circumstances.

The distinction of cell cycle states between the endothelial cells of veins and arteries is maintained into adulthood. Thus, flow-mediated regulation of endothelial cell cycle state may be broadly required for adult vascular homeostasis and its dysregulation may contribute to vascular diseases. Indeed, arterial-venous malformations can be associated with underlying endothelial cell hyperproliferation that may contribute to disruptions in arterial-venous identity^17,27-29^. This is of particular interest given that pharmacological cell cycle arrest in late G1 was found to be sufficient to rescue arterial-venous specification and maturation in an animal model of dysregulated vascular network formation. These experiments employed the CDK4/6 inhibitor palbocilib that is FDA approved for the treatment of HR-positive, HER2-negative metastatic breast cancer^30,31^, and is currently being investigated for other forms of cancer^32,33,34^. Thus, palbocilib or similar drugs may be beneficial for the treatment of vascular diseases, such as arteriovenous and cerebral cavernous malformations, and they may also have utility for the control arterial-venous identity during the creation of arteriovenous fistulas, coronary artery bypass grafting and vascular procedures.

## Acknowledgements

We thank the scientists at the Yale Center for Genomic Analysis, Yale Flow Cytometry Facility, and the University of Virginia Flow Cytometry Facility for their expert technical ability and advice. We also thank the University of Virginia Health Sciences Library and Research Computing centers for their expert advice and guidance on bioinformatic analysis. We greatly appreciate the lab of Dr. Martin Schwartz (Professor, Yale University) for providing equipment and technical advice on shear flow experiments, and the lab of Dr. Patrick Gallagher (Professor, Yale University) for providing advice and reagents for the ATAC-sequencing experiments. The experiments in this study were supported by NIH grants to N.W.C. (T32 HL007224, T32 HL007284), K.W. (R01 HL142650 and R01 HL141256) and K.K.H. (R01 HL146056 and U2EB017103), and AHA grant to N.G. (19POST34400065).

## Author contributions

N.W.C. coordinated the project and wrote the manuscript. N.W.C., G.G., M.P., N.G., C.M., K.W., and K.K.H. contributed to experimental design and data analysis. N.W.C. performed retina cell cycle analysis, next-generation sequencing and bioinformatics analysis, and *in vitro* mechanistic studies. G.G. performed *in vitro* protein analysis. G.G. and M.P. performed *in vivo* CDK4/6i experiments. N.G. and C.M. assisted with image analysis and supplemental control experiments. H.V. and E.N. generated HUVEC-FUCCI cell lines. A.K. assisted with bioinformatics analysis. S.M. and M.H. maintained mouse lines and performed *in vivo* experiments.

## Materials and Methods

### Mouse strains

All animal procedures were approved by the Institute for Animal Care and Use Committees at Yale University and the University of Virginia. In this study, FUCCI-26aP mice^7^ were bred with Cdh5-CreER^T2^ mice^14^ to generate endothelial-specific FUCCI expression after tamoxifen injection. Tamoxifen (Sigma Cat# T5648) was resuspended in 10% EtOH and 90% Corn oil (Sigma Cat# C8267) at 4 mg/mL, and 25 μL was injected per pup. Additionally, Gja4-/- (Cx37-KO) mice^35^ were bred with R26p-Fucci2 mice^36^ to generate FUCCI expression in Cx37-KO mice.

### Retina isolation, staining and quantification

Retinas were isolated from mice, stained and imaged as previously described^4^. All antibodies are listed in Table S1. For retina imaging, isolated retinas were immunostained for CD31, Erg1/2/3, or αSMA, with fluorescent secondary antibodies. Stained retinas were imaged by confocal microscopy (Leica SP8 or Leica SP5). Endothelial cells along arterial branches and venous branches were quantified by Erg1/2/3 expression in the nuclei, and cell cycle was determined by FUCCI expression (Early G1 by mCherry^-^mVenus^-^; Late G1 by mCherry^+^mVenus^-^; S/G2/M by mCherry^-^mVenus^+^). For isolation of endothelial cells from retinas, dissected retinas were digested with Collagenase Type II (1.0 mg/mL, Gibco Cat# 17101015) in DMEM (Gibco Cat# 21013024) and 10% FBS (Gibco Cat# 26140079) for 20 min, washed, stained for anti-CD31 and -CD45 in staining buffer (HBSS with 10% FBS, 20 mM HEPES, 1 mg/mL D-Glucose), then resuspended in FACS buffer (PBS with 1% FBS). FUCCI cell populations were isolated by FACS through a CD31^+^CD45^-^ gating strategy, then by mCherry/mVenus to determine cell cycle. Cells were sorted into RNA lysis buffer, and RNA was purified with RNeasy Micro Kit (Qiagen Cat# 74034). FACS was performed with a BD FACSAria at either the Yale Flow Cytometry Core or the University of Virginia Flow Cytometry Core.

### Quantitative RT-PCR

Purified RNA was converted to cDNA using the High-Capacity cDNA Reverse Transcription Kit (ThermoFisher Cat# 4368814) and quantified by Power SYBR Green PCR Master Mix (ThermoFisher Cat# 4368577) via qRT-PCR (Applied Biosystems QuantStudio 6). Gene-specific primers are listed in Table S2. Relative quantification was determined by the delta-delta-CT method.

### Generation of HUVEC-FUCCI

Primary Human Umbilical Vein Endothelial Cells (HUVEC) were obtained from either the Yale Vascular Biology and Therapeutics Core. HUVEC were passaged in Endothelial Cell Growth Medium (PromoCell Cat# C-22010) and experiments were performed in EGM-2 (Lonza Cat# CC-3162). Experiments used cells between passage 4 and 8. HUVEC were infected with lentivirus generated with HEK293T cells infected with the Fast-FUCCI plasmid^13^; pBOB-EF1-FastFUCCI-Puro was a gift from Kevin Brindle & Duncan Jodrell (Addgene plasmid # 86849; http://n2t.net/addgene:86849; RRID:Addgene_86849). Cells were selected and passaged in Puromycin (1 μg/mL, Sigma P9620).

### HUVEC-FUCCI treatment with shear stress and ligands

HUVEC-FUCCI were plated on 25-55mm cell plastic slides pre-treated with 10 μg/mL fibronectin, then subjected to venous (4 dynes/cm^2^) or arterial (12 dynes/cm^2^) shear flow forces in parallel plate flow chambers (pressure-dampened, gravity-driven dual reservoir system with media re-circulating via a peristaltic pump (Masterflex, Cole Palmer) and maintained at 37 C, 5% CO2) for 24 hr, as previously described^37^. Additionally, HUVEC-FUCCI were seeded in a 6-well plate at 5×10^4^ cells per well, then treated with control media or media supplemented with TGF-β1 (1 ng/mL, R&D Systems Cat# 240-B) or BMP4 (5 ng/mL (?), R&D Systems Cat# 314-BP). Cells were incubated with ligand for 8 hours. Cells were then lifted by Trypsin, and cell cycle was determined by flow cytometry with FUCCI reporter expression.

### FACS of HUVEC-FUCCI

Sorting of HUVEC-FUCCI into cell cycle states was performed by lifting subconfluent cells from cell culture with Accutase (Sigma Cat# A6964), washing cells and resuspending in FACS buffer. Fluorescent levels of mCherry and mVenus were used to determine cell cycle state. Cells were either sorted by FACS with a BD FACSAria or BD FACSMelody, or analyzed by flow cytometry with a BD LSRII.

### HUVEC-FUCCI bulk RNA sequencing and data analysis

RNA from HUVEC-FUCCI sorted into different cell cycle states was purified and submitted for next-generation transcriptome sequencing to the Yale Center for Genomic Analysis (Illumina HiSeq4000). Raw read data was quality-control checked (FastQC), aligned to human genome GRCh38 using Kallisto^38^, and analyzed for total and differential gene expression using Sleuth^39^ (Supplemental Data 1). Gene Ontology analysis was performed with GAGE^40^ (Supplemental Data 2).

### Western blot analysis

Protein was isolated from cells using RIPA Buffer (Sigma Cat# R0278) or sorted directly into Laemmli Buffer (BioRad Cat# 1610747). Western blot analysis was performed using the Criterion Vertical Electrophoresis Cell (BioRad Cat# 1656020) with 4%-15% Criterion Tris-HCl Protein Gels (BioRad Cat# 3450028) and imaged with the Azure Biosystems c300. Western blots were quantified by ImageJ densitometry analysis.

### TGF-β1/BMP4 ligand induction

Subconfluent HUVEC-FUCCI were lifted and sorted into early G1 and late G1 cell cycle states, then seeded at 5×10^4^ cells per well into 6-well plates. Cells were left to attach to the plate for 1 hr in a 37° C, 5% CO_2_ incubator, then media was changed with new media supplemented with control, TGF-β1 (1 ng/mL, R&D Systems Cat# 240-B) or BMP4 (5 ng/mL, R&D Systems Cat# 314-BP). For co-immunoprecipitation experiments, cells were incubated in ligands for 2 hr, then SMAD4 complexes were isolated using the Pierce Co-Immunoprecipitation Kit (ThermoFisher Cat# 26149) per manufacturer’s instructions. For chromatin immunoprecipitation experiments, cells were incubated in ligands for 4 hr, then SMAD4-DNA complexes were isolated using the High-Sensitivity ChIP Kit (AbCam Cat# ab185913), per manufacturer’s instructions. For gene induction experiments, cells were incubated in ligands for 8 hr, then RNA lysate was collected and qRT-PCR was performed.

### HUVEC-FUCCI ATAC-sequencing and data analysis

Subconfluent HUVEC-FUCCI were lifted and sorted into early G1 and late G1 cell cycle states, then immediately collected for ATAC-sequencing analysis. Library preparation was performed as previously described^41^. Sequencing was performed at the Yale Center for Genomic Analysis (Illumina HiSeq4000). Raw read data was quality controlled with FastQC (Babraham Bioinformatics), filtered and trimmed with Trimmomatic^42^, then peaks were called with MACS2^43^, and differential peak analysis was performed with HOMER (UCSD) (Supplemental Data 3).

### Transfection of siRNA

HUVEC-FUCCI were transfected with ThermoFisher Silencer Select siRNA targeting SMAD1 (Cat# s8394), SMAD2 (Cat# s8397), SMAD3 (Cat# s8400), or SMAD5 (Cat# s8406). RNAiMAX Lipofectamine (ThermoFisher Cat# 13778075) was used to package and transfect siRNA into HUVEC-FUCCI, per manufacturer’s instructions. After 48 hr, HUVEC-FUCCI were lifted and sorted by FACS into early G1 or late G1, then induced with ligand, as previously described.

### Cdk4/6 inhibitor administration in mice

Cdk4/6 inhibition in mice was performed by resuspending Palbociclib (Sigma-Aldrich Cat# PZ0383) in 50 mM Sodium lactate (Sigma Cat# L7022) at 12 mg/mL, then administered to pups at P3, P4, and P5 by oral gavage. Retinas were isolated at P6. EdU incorporation assay was performed using the Click-It EdU Cell Proliferation Kit (ThermoFisher Cat# C10340) per manufacturer’s instructions, with EdU injection into mice 5 hours before euthanasia at P6.

### Statistical analysis

Unless otherwise indicated, statistical analysis was performed using either a standard two-tail Student’s t-test or a two-way ANOVA test followed by a Tukey’s multiple comparison corrected post-hoc test. All statistical analysis of RNA sequencing and ATAC sequencing data sets was performed through computational analysis packages, which contain statistical corrections for large data sets.

**Table S1.** Antibodies used in immunofluorescence, FACS, western blot and immunoprecipitation

| **Application** | **Antibody** | **Source** |
| --- | --- | --- |
| Immunofluorescence | Goat anti-Mouse CD31 | R&D Systems Cat# AF3682 |
|  | Rabbit anti-Mouse ERG1/2/3 | AbCam Cat# ab92513 |
| | Mouse anti-Mouse αSMA | ThermoFisher Cat# 50-9760-82 |
| FACS | CD31-APC Rat anti-Mouse | BD Biosciences Cat# 551262 |
|  | CD45-V450 Rag anti-Mouse | BD Biosciences Cat# 560501 |
| Western Blot | Goat anti-TGFBR1 | R&D Systems Cat# AF3025 |
|  | Phospho-SMAD3 Rabbit mAb | Cell Signaling Cat# 9520 |
|  | Phospho-SMAD1/5/9 Rabbit mAb | Cell Signaling Cat# 13820 |
|  | B Actin Rabbit mAb | Cell Signaling Cat# 4970 |
|  | SMAD2/3 Rabbit mAb | Cell Signaling Cat# 8685 |
|  | SMAD1 Rabbit mAb | Cell Signaling Cat# 6944 |
|  | SMAD5 Rabbit mAb | Cell Signaling Cat# 12534 |
|  | SMAD4 Rabbit mAb | Cell Signaling Cat# 46535 |
|  | Rabbit anti-BMPR2 | AbCam Cat# ab96826 |
|  | Goat anti-Human ALK1 | R&D Systems Cat# AF370 |
|  | Rabbit anti-TGF beta RII | AbCam Cat# ab186838 |
|  | Goat anti-Human Endoglin | R&D Systems Cat# AF1097 |
|  | Phospho p44/42 MAPK (Erk1/2) | Cell Signaling Cat# 9106 |
|  | P44/42 MAPK (Erk1/2) | Cell Signaling Cat# 9102 |
|  | Akt Rabbit Ab | Cell Signaling Cat# 9272 |
|  | Phospho Akt (Ser473) Rabbit mAb | Cell Signaling Cat# 4060 |
|  | Horse anti-Goat IgG (H+L) | Vector Labs Cat# PI-9500 |
|  | Goat anti-Rabbit IgG (H+L) | Vector Labs Cat# PI-1000 |
| Immunoprecipitation | SMAD4 Rabbit mAb | Cell Signaling Cat# 46535 |

**Table S2.**
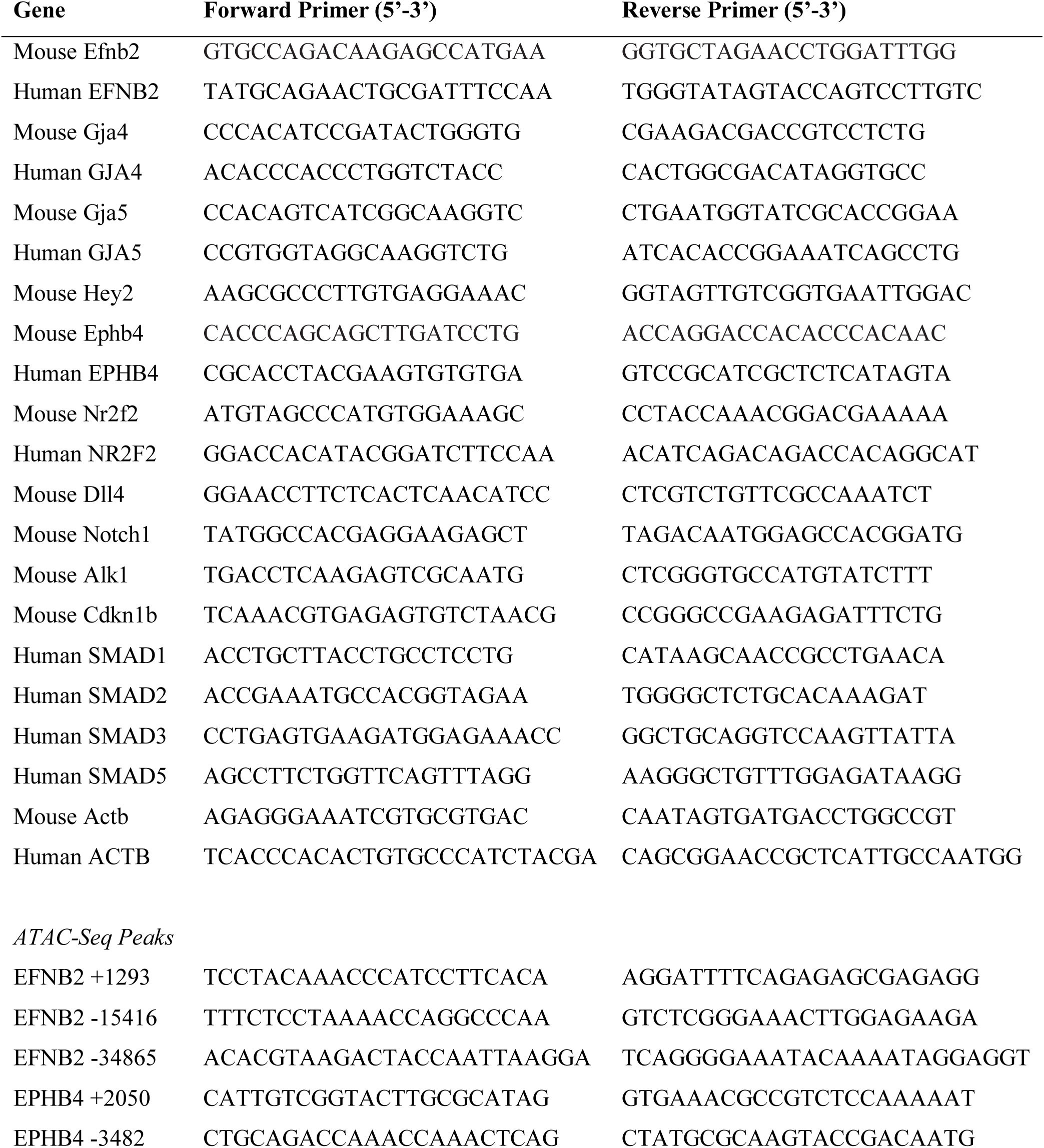
Gene-specific primers for qRT-PCR

**Supplemental Data 1**. Gene expression results from bulk RNA sequencing of early G1 and late G1 HUVEC-FUCCI

*Additional attachment*

**Supplemental Data 2**. Gene ontology results from bulk RNA sequencing of early G1 and late G1 HUVEC-FUCCI

*Additional attachment*

**Supplemental Data 3**. Peak quantification results from ATAC sequencing of early G1 and late G1 HUVEC-FUCCI

*Additional attachment*

## Figure legends

**Extended Figure 1.**
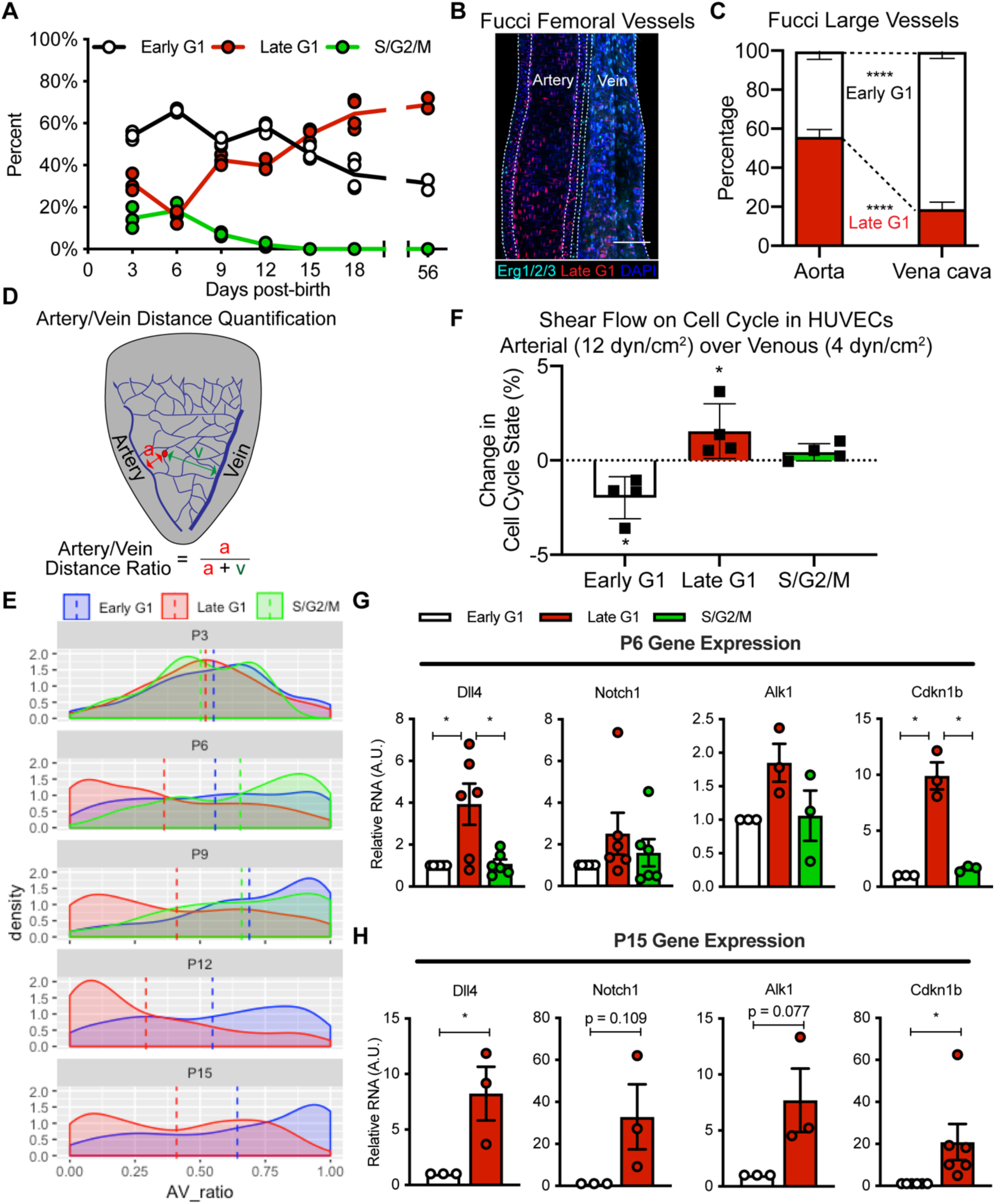
Endothelial Cell Cycle State During Retina Vascular Development. **A**) Cell cycle state of FUCCI2 mouse retinal endothelial cells. Endothelial cell cycle state in **B**) confocal z-stack imaged femoral vessels, and **C**) quantified aorta and vena cava endothelial cells. **D**) Overview of Artery/Vein Distance Ratio determination. **E**) Probability density of Artery/Vein Distance Ratio for retinal endothelial cells in early G1, late G1 or S/G2/M. **F**) HUVEC-FUCCI cell cycle changes in response to arterial and venous shear stress. Gene expression of Dll4, Notch1, Alk1, and Cdkn1b in retinal endothelial cells in cell cycle states at **G**) P6 and **H**) P15.

**Extended Figure 2.**
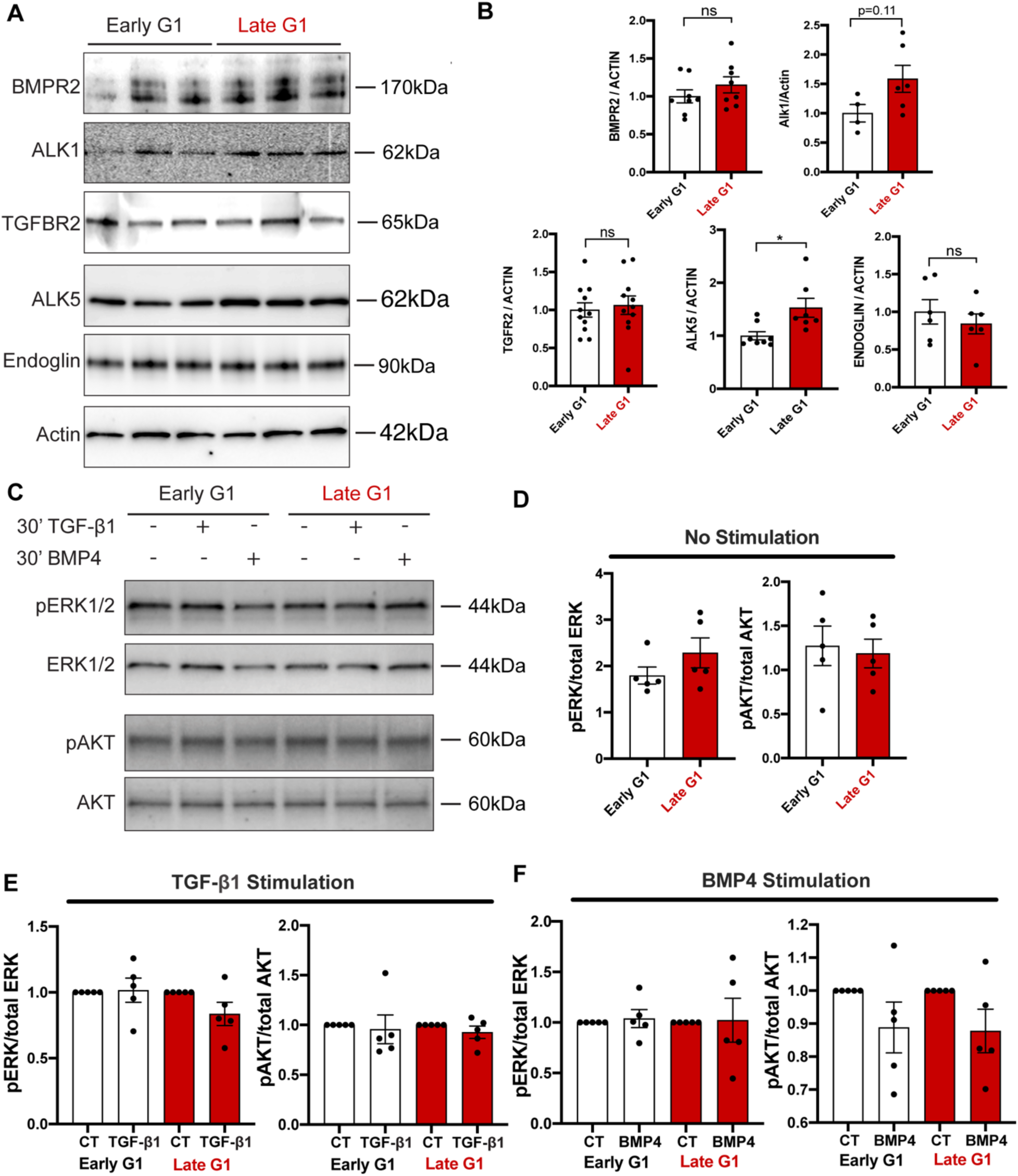
Cell Cycle-Dependent Expression of TGF-β/BMP Pathway in HUVEC-FUCCI. **A**) Western blot of TGF-β/BMP signaling proteins in HUVEC-FUCCI in early G1 and late G1 (n = 3), **B**) quantified. **C**) Western blot of ERK1/2 and AKT phosphorylation in HUVEC-FUCCI in early G1 and late G1 after TGF-β1 or BMP4 treatment, **D-F**) quantified.

**Extended Figure 3.**
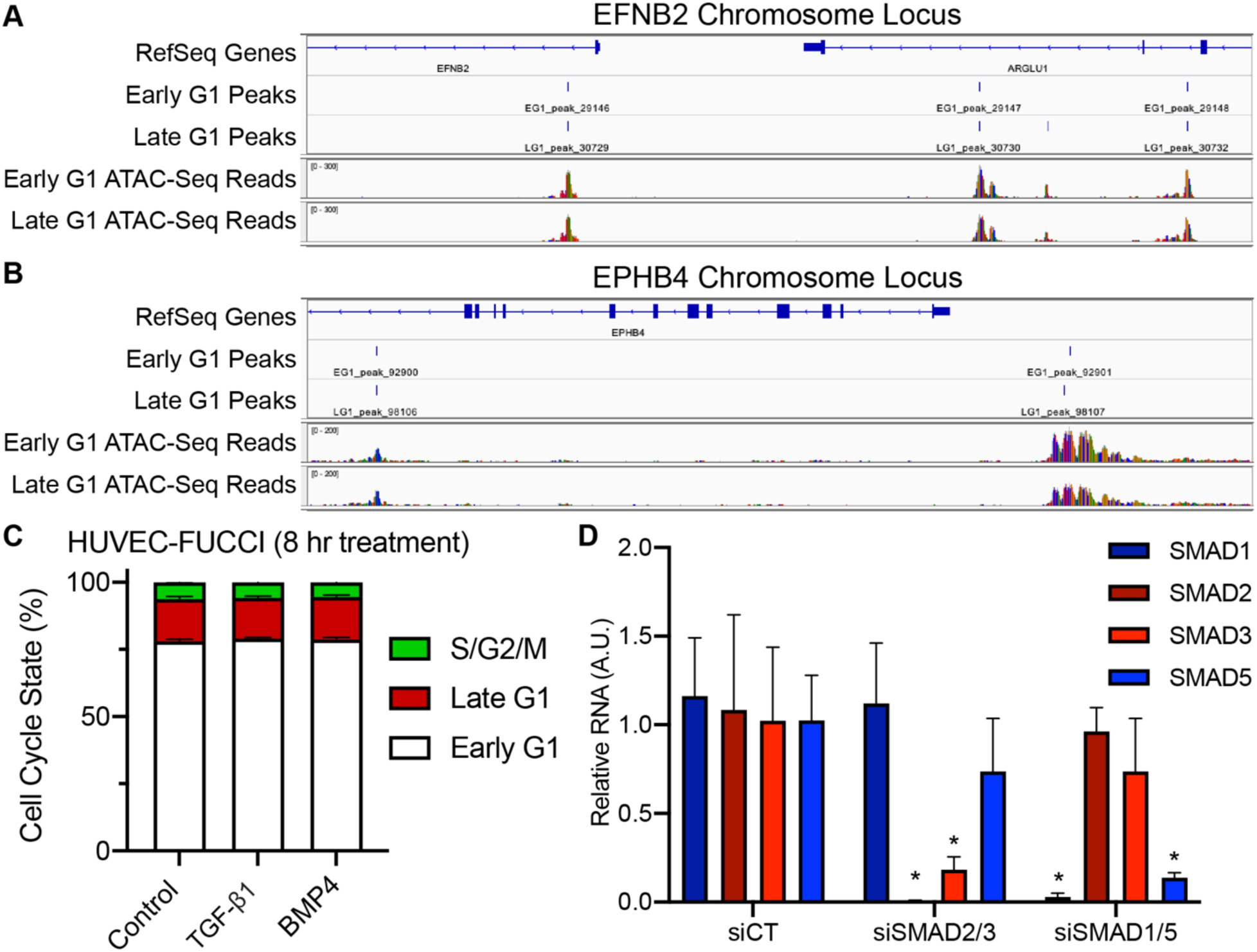
Cell Cycle-Dependent Arterial-venous Specification via TGF-β/BMP Signaling in HUVEC-FUCCI. Peaks from ATAC-Sequencing of HUVEC-FUCCI in early G1 and late G1 around the **A**) EFNB2 locus, and **B**) EPHB4 locus. **C**) HUVEC-FUCCI cell cycle states after TGF-β1 or BMP4 treatment. **D**) qRT-PCR of SMAD genes in HUVEC-FUCCI after SMAD siRNA transfection.

**Extended Figure 4.**
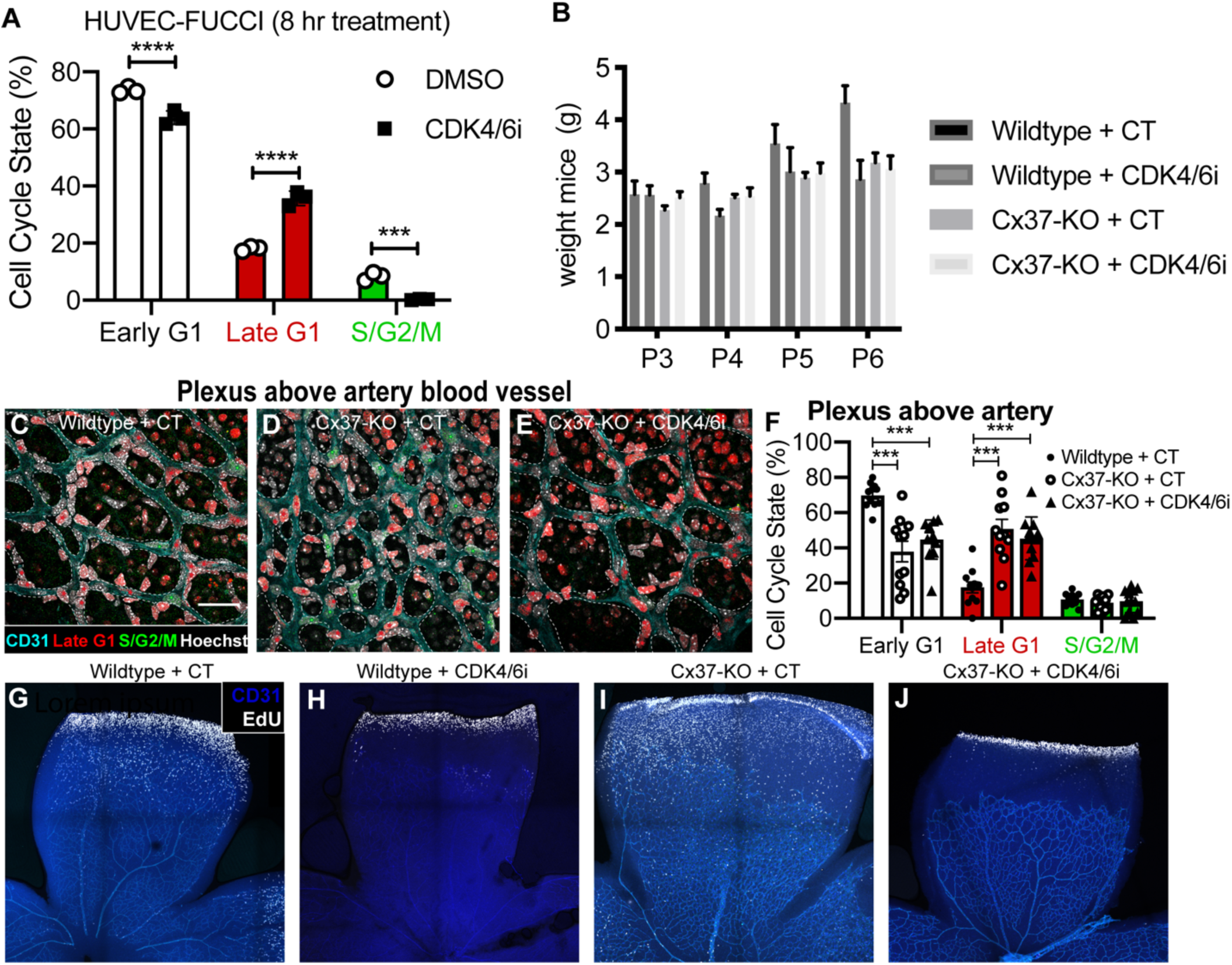
Rescue of Arterial-vvenous Specification Defects with Pharmacological CDK4/6 Inhibition. **A**) HUVEC-FUCCI cell cycle states after CDK4/6i treatment. **B**) WT, WT+CDK4/6i, Cx37-KO and Cx37-KO+CDK4/6i treated mice analyzed for weight over time. **C-F**) P6 retinal vasculature of FUCCI2, FUCCI2+Cx37-KO and FUCCI2+Cx37-KO+CDK4/6i treated mice imaged for CD31, hCdt1(30/120), hGem(1/110) and Erg1/2/3 and quantified for cell cycle state in plexi above arterial blood vessels (scale bars = 50μm, vessels outlined in dotted white lines, cell cycle state highlighted with colored stars). **G-J**) WT, WT+CDK4/6i, Cx37-KO and Cx37-KO+CDK4/6i treated mice analyzed for EdU incorporation.

## Notes

### Competing Interest Statement

The authors have declared no competing interest.

